# Optogenetic ion pumps differ with respect to the secondary pattern of K^+^ redistribution

**DOI:** 10.1101/2023.04.13.536767

**Authors:** R. Ryley Parrish, Tom Jackson-Taylor, Juha Voipio, Andrew J. Trevelyan

**Affiliations:** Newcastle University Biosciences Institute, Medical School, Framlington Place, Newcastle upon Tyne, NE2 4HH, UK; Department of Cell Biology and Physiology, Brigham Young University, Provo, Utah. 84602, USA; Faculty of Biological and Environmental Sciences, Molecular and Integrative Biosciences, University of Helsinki, 00014 Helsinki, Finland

**Keywords:** Archaerhodopsin, Halorhodopsin, chloride-cation-cotransporter, potassium, chloride

## Abstract

We recently reported that strong activation of the optogenetic chloride pump, Halorhodopsin leads to a secondary redistribution of K^+^ ions into the cell, through tonically open, “leak” K^+^ channels. Here we show that this effect is not unique to halorhodopsin, but is also seen with activation of another electrogenic ion pump, archaerhodopsin. The two opsins differ however in the size of the rebound rise in extracellular potassium, [K^+^]o, after the end of activation, which is far larger with halorhodopsin than for archaerhodopsin activation. Multiple linear regression modelling indicates that most of the variance in the post-illumination surge in [K^+^]o was explained by the type of opsin, and almost nothing by the size of the preceding, illumination-induced drop in [K^+^]o. These data provide additional support for the hypothesis that intense chloride-loading of cells, as occurs naturally following intense bursts of GABAergic synaptic bombardment, or artificially following halorhodopsin activation, is followed by extrusion of both Cl and K^+^ coupled together. We discuss this with respect to the pattern of [K^+^]o rise that occurs at the onset of seizure-like events.

## Introduction

The development of optogenetics techniques and tools have presented new ways of manipulating neuronal excitability (Deisseroth, 2015), using either light activated ion channels (Boyden et al., 2005) or electrogenic ion pumps (Chow et al., 2010; Zhang et al., 2007). The real power of these remarkable tools is that they may be targeted very precisely to specific subsets of cells, or even to subcellular domains, allowing experimental control to be delivered at these sites. The induced ion movements, however, do alter the internal and external milieu, meaning that there may be unintended secondary effects both at and away from the site of expression. These, though, may be put to good use for exploring neuronal physiology and pathology, so it is worth documenting these effects.

Halorhodopsin was the first optogenetic protein to be used for inhibiting neurons, using light to drive chloride into neurons to hyperpolarize them (Zhang et al., 2007). We recently showed that this light-activated chloride movement has several interesting secondary effects (Parrish et al., 2023). Firstly, the hyperpolarization of membrane potential alters the balance between the electrical and chemical gradients for other ions; since the open “leak” channels at rest are mainly K^+^ channels, there follows a redistribution of K^+^ ions into expressing cells, causing the extracellular K^+^ concentration, [K^+^]_o_, to drop. Consequently, there is a negative shift in membrane potential in other non-expressing neurons, with a measurable drop in their excitability (an “off-target” inhibition that may be important to recognize when interpreting optogenetic experiments). Most surprising, though, was that activation of halorhodopsin triggered cortical spreading depolarization (CSD) events that occurred even during periods of ongoing illumination, when most neurons were hyperpolarized and [K^+^]_o_ was below baseline levels. Interestingly, on the trials where CSD were not induced and once illumination ended, there was then a large rebound surge in [K^+^]_o_.

Archaerhodopsin, another commonly used inhibitory optogenetic ion pump, delivers hyperpolarization of neurons in a different way, by driving protons out of the cell (Chow et al., 2010). As such, archaerhodopsin activation should show some of the secondary effects of halorhodopsin, but not others. It has already been shown that activation of halorhodopsin, but not archaerhodopin, induces a pronounced chloride-loading of neurons that may persist for many seconds (Alfonsa et al., 2015; Raimondo et al., 2012). Since Cl^-^ clearance is achieved primarily by coupling to K^+^ extrusion, through the electroneutral chloride-cation cotransporter, KCC2 (Kaila et al., 2014; Rivera et al., 1999), the degree of chloride loading will influence the amplitude of the post-illumination surge in [K^+^]_o_. To examine these differences, we present recordings of [K^+^]_o_ in neocortical brain slices in which archaerhodopsin is expressed widely in the pyramidal population, to show that it shares with halorhodopsin the effect of inducing an inward movement of K^+^ ions during the period of illumination. In contrast, the post-illumination rebound surge in [K^+^]_o_ following archaerhodopsin activation appears much smaller than for halorhodopsin.

## Methods

### Ethical Approval

All procedures performed were in accordance with the guidelines of the Home Office UK and Animals (Scientific Procedures) Act 1986 and approved by the Animal Welfare and Ethical Review Body at both Newcastle University and UCL. Male and female mice were used for experimentation.

### Archaerhodopsin viral injections

C57/B6 adult mice were injected with AAV9.CAG.ArchT, purchased from the UPenn vector core. For these adult injections, done at 8-12 weeks of age, animals were anesthetized by ketamine–methoxamine intraperitoneal injection and placed in a stereotaxic head holder (David Kopf Instruments). Injections were made at three to four locations in an anterior–posterior row in one hemisphere, 1.5–2 mm lateral to the midline and 1–0.4 mm deep to the pia (0.6 μl, injected over 15 min).

### Slice Preparation

We used both wild-type C57/B6 mice, and also mice expressing eNpHR3.0 within the pyramidal cell population, generated by cross-breeding homozygous Emx1-cre mice (Jackson Laboratory, Stock 005628) with mice containing a floxed STOP cassette in front of an eNpHR3.0/EYFP domain (Jackson Laboratory, Stock 014539; both maintained on the C57/B6 background). Emx1-promoter yields gene expression only in pyramidal cells in neocortex, in adult mice, although it can drive gene in glia in other brain areas, and early in development (Gorski et al., 2002). Experiments were performed on mice aged 1 - 8 months, of both sexes. Mice were first decapitated, brains were removed and placed in cold cutting solution containing (in mM): 3 MgCl2; 126 NaCl; 2.6 NaHCO3; 3.5 KCl; 1.26 NaH2PO4; 10 glucose. 400 µm horizontal sections were made on a Leica VT1200 vibratome (Leica Microsystem, Germany). Slices were stored at room temperature, in an interface holding chamber for 1 – 4 hours prior to experimentation. Solutions were bubbled with carboxygen (95% O2 and 5% CO2) in artificial cerebro-spinal fluid (aCSF) containing (in mM): 2 CaCl2; 1 MgCl2; 126 NaCl; 26 NaHCO3; 3.5 KCl; 1.26 NaH2PO4; 10 glucose.

### *In vitro* Extracellular recordings

Extracellular recordings were performed using an interface recording chamber. Slices were placed in the recording chamber perfused with either aCSF, supplemented in different experiments with different combinations of the following drugs: 1µM Tetrodotoxin (TTX) (Abcam), 10mM tetraethylammonium (TEA) (Sigma), as indicated in the results section. Recordings were obtained using aCSF-filled ∼1-3MΩ borosilicate glass microelectrodes (GC120TF-10; Harvard apparatus, Kent) placed in deep layers of neocortex. Extracellular potassium [K^+^]_o_ was measured using single-barrelled K^+^-selective microelectrodes. The pipettes were pulled from nonfilamented borosilicate glass (Harvard Apparatus, Kent, UK), and the glass was exposed to vapor of dimethyl-trimethyl-silylamine (Sigma-Aldrich), baking at 200^0^ C for 40 minutes. The pipettes were then backfilled with aCSF. A short column of the K^+^ sensor (Potassium ionophore I, cocktail B; Sigma-Aldrich, #99373) was taken into the tip of the salinized pipette by using slight suction. The recordings through the K^+^-sensor electrode were referenced to a second electrode filled with aCSF. From the differential signal from a custom build amplifier, we calculated the [K^+^]_o_ from calibration recordings made in an open bath, using sudden increments in [K^+^]_o_. This provided a scaling factor S, of 55-59mV, where the K^+^ concentration at a given moment in time, t, was calculated from the differential voltage, V(t), as follows: [K]_o_ = [K]_o.baseline_ 10^V(t)/S^ [K]_o.baseline_ for our experiments was 3.5 mM. The temperature of the chamber and perfusate was maintained at 33-36 C using a closed circulating heater (FH16D, Grant instruments, Cambridge, UK). The solutions were perfused at the rate of 3 mls/min by a Watson Marlow 501U peristaltic pump (Watson-Marlow Pumps Limited, Cornwall UK). For the collection of the HaloRhodopsin and ArchT data, the direct current local field potential signal was unfiltered and amplified to a 10X output with a custom build amplifier. These waveform signals were digitized with a Micro 1401-3 ADC board (Cambridge Electronic Design, UK) and Spike2 version 7.10 software (Cambridge Electronic Design, UK). Signals were sampled at 10 kHz. Recordings were analysed using a custom-written code in Matlab2015b (Mathworks, MA, USA).

### Optogenetic illumination for extracellular recordings

Optogenetic illumination was delivered for 90s continuous periods, using a Fiber-coupled LED light at 565 nm (Thorlabs, M565F3) and driven by an LED driver (Thorlabs, LEDD1) placed in trigger mode. Light output was measured at an average of 9 mW. The LED light was positioned just above the superficial layers of the neocortex for the slice experiments and pointed toward the deep layers, where the extracellular recordings were performed.

Statistics: Statistical analysis of electrophysiology was performed using GraphPad Prism (GraphPad Software, Inc., La Jolla, CA, USA). Data was analysed using Mann-Whitney test (MATLAB, *ranksum*.*m*) or multiple linear regression modelling (*fitlm*.*m*). Figures of electrophysiology traces were created in Matlab2015b or 2018b (Mathworks, MA, USA).

## Results

### Activation of archaerhodopsin causes secondary redistribution of K^+^ ions

We previously reported that activation of the optogenetic chloride pump, halorhodopsin, induced a secondary ionic redistribution of K^+^ ions into the cell during the period of activation, followed by a surge in [K^+^]_o_ after illumination ended (Parrish et al., 2023). The inward movement of K^+^ appears to pass through leak K^+^ channels, whereas the post-illumination outward movement included a KCC2-sensitive component, implying that it was driven in part by the need to remove Cl^-^ ions that had been pumped into cells. We reasoned therefore that these effects may be replicated in part by activation of other optogenetic pumps. To test this prediction, we recorded the pattern of K^+^ redistribution associated with activation of the proton pump, archaerhodopsin (Figure 1). Widespread archaerhodopsin expression in the pyramidal cell population was achieved by injecting a viral vector carrying the floxed gene into mice expressing Cre-recombinase under the Emx1 promoter (Gorski et al., 2002). During periods of illumination, we recorded a highly significant drop in [K^+^]_o_ in brain slices expressing archaerhodopsin (Figure 1), but not in brain slices from wild-type mice. This K^+^ redistribution was almost entirely eliminated by blockade of plasmalemmal K^+^ channels, using TEA. In both respects, archaerhodopsin mimicked the effect we had previously reported in halorhodopsin-expressing brain slices. In our preparations, the amplitude of the K^+^ redistribution was larger in the halorhodopsin-expressing brain slices, which might reflect the degree of hyperpolarization achieved.

**Figure 1.**
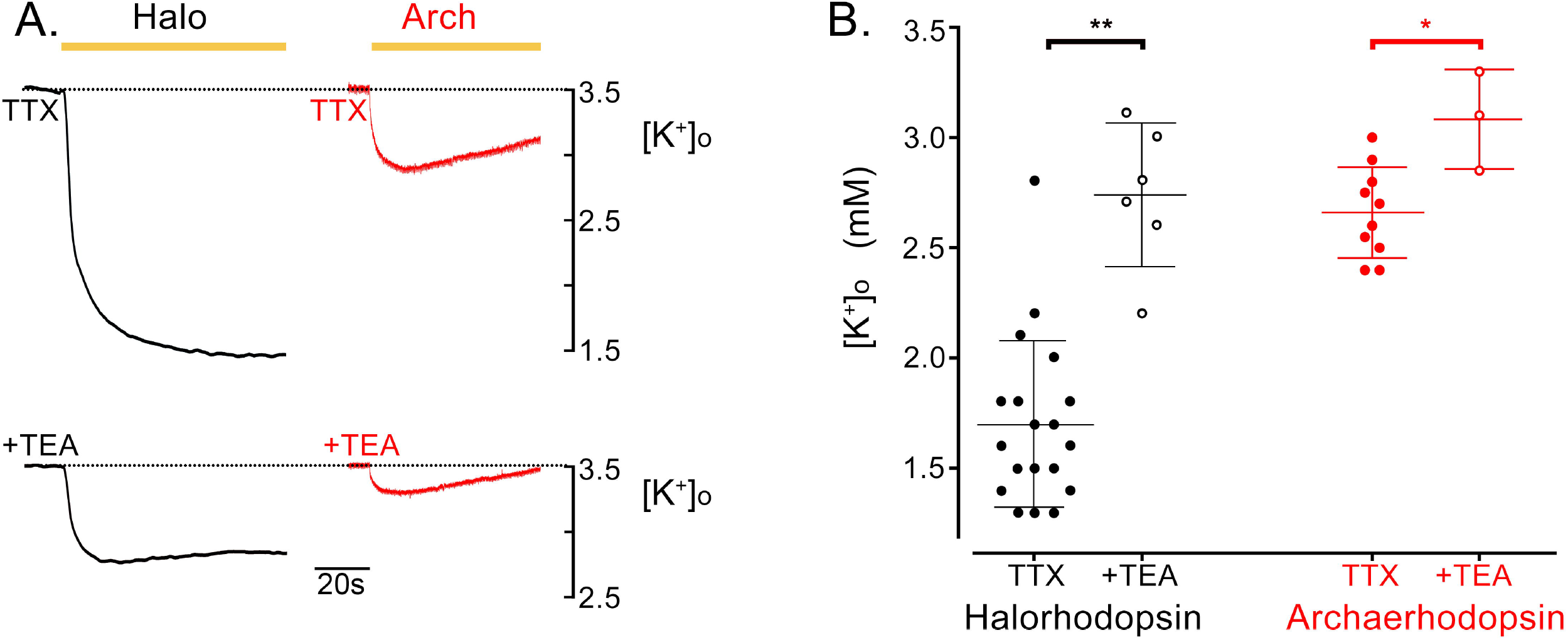
The main movement of K^+^ ions from the extracellular space, induced by opsin activation is through K^+^ channels. (A) Redistribution of K^+^, assessed by recording [K^+^]_o_ during a period of halorhodopsin (black) or of archaerhodopsin activation (red) in the presence of TTX to block neuronal firing, and additionally TEA to block “leak” K^+^ channels. (B) Pooled data of measurements of [K^+^]_o_ for all the halorhodopsin and archaerhodopsin recordings. For both opsins, the size of the K^+^ redistribution is greatly reduced by block of voltage-gated K^+^ channels (Halo vs Halo + TEA, p = 0.00076; Arch vs Arch + TEA, p = 0.0279, two-tailed Mann-Whitney test).

At the end of the illumination period, we previously reported that halorhodopsin-expressing brain slices showed a very large rebound surge in [K^+^]_o_ (7.98 ± 0.41 (mean ± s.e.m), range 5.5 - 10.6 mM; 95% confidence interval = 7.18 - 8.78 mM). In contrast, the post-illumination surge in archaerhodopsin-expressing slices was significantly above the 3.5mM baseline level (4.12 ± 0.12 mM, range 3.4 - 5.1 mM; 95% confidence interval = 3.86 - 4.38 mM), but significantly smaller than occurred with the halorhodopsin slices (ArchT vs Halorhodopsin, p = 0.0001, two-tailed Mann-Whitey test) (Figure 2).

**Figure 2.**
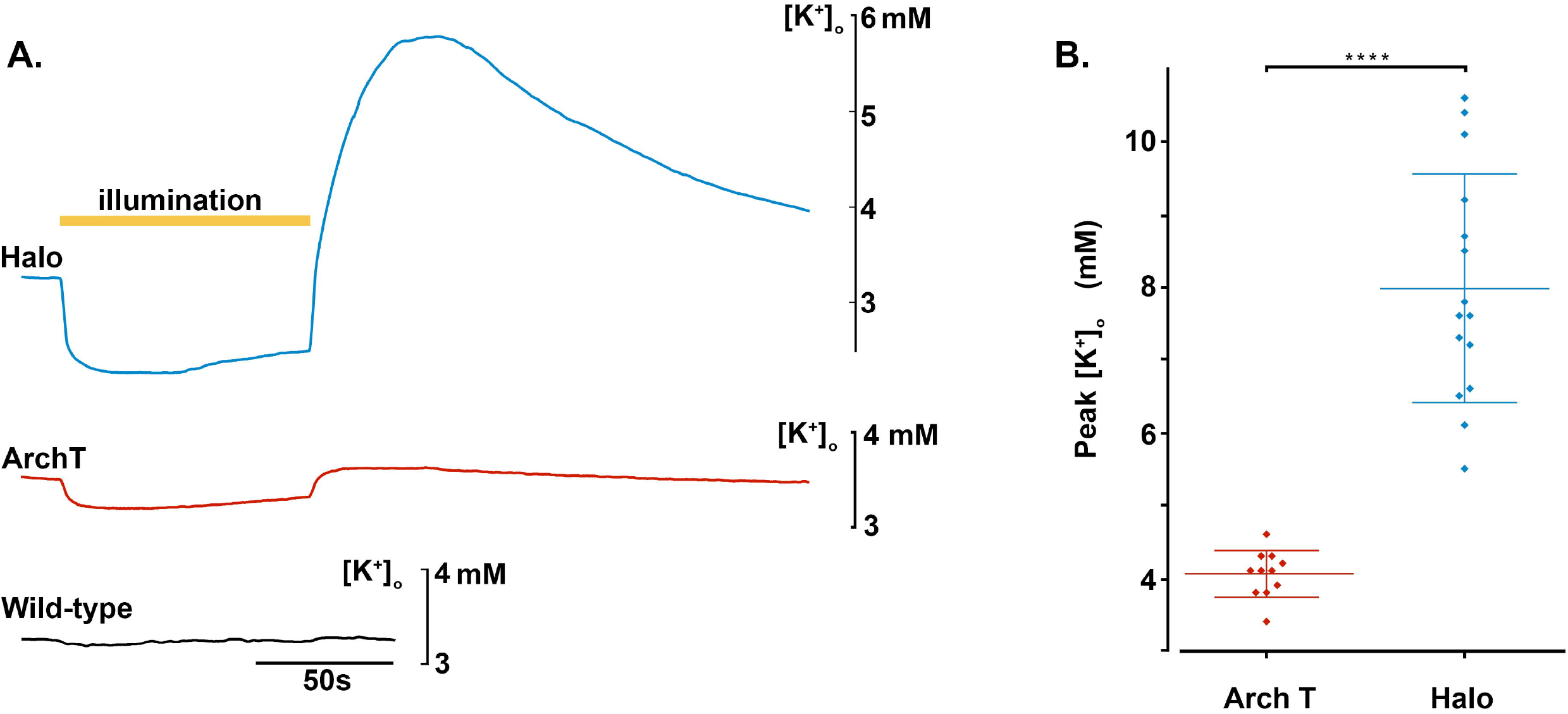
The large post-illumination surge in [K^+^]o following a period of halorhodopsin activation is not replicated by archaerhodopsin activation. (A) Recordings of extracellular [K^+^] in brain slices in which either Halorhodopsin (blue, top), or Archaerhodopsin (red, middle), was expressed in all pyramidal cells, or in a slice with no opsin expression (black, bottom). Yellow bar represents the time of illumination (90s period). Light activation of Halorhodopsin induced a very marked reduction in [K^+^]_o_, followed by an even larger, rebound increase. There was also a drop in [K^+^]_o_, synchronous with activation of Archaerhodopsin, but the rebound was negligible. Brain slices without opsins showed no light induced fluctuations in [K^+^]_o_. (B) The peak extracellular [K^+^], after the end of illumination, for Archaerhodopsin (mean peak = 4.12mM, n = 11) and Halorhodopsin activation (mean peak = 7.98mM; n = 15; comparison, p = 0.0001, two-tailed Mann-Whitey test), respectively.

For both opsins, a component of the rebound surge will be extrusion of K^+^ ions that moved into the neurons during the period of illumination; since the illumination-induced inward movement was greater in our halorhodopsin group (possibly due to stronger expression; during Halorhodopsin activation, the range of the drop in of drop in [K^+^]_o_ = 1.2 - 2.2 mM below baseline (3.5mM); during ArchT activation, the range = 0.5 – 1.3 mM below baseline), one possibility is that the larger rebound in that group simply reflected this difference. One approach to test this would have been to adjust expression patterns or illumination levels to try to match the amplitude of the illumination-induced drop for the two different opsins, but the halorhodopsin experiments were performed first, and we were not able to achieve the same degree of K^+^ redistribution with the archaerhodopsin. Instead, we pooled the datasets and used multilinear regression modelling of the [K^+^]_o_, with the predictive variables being the illumination-induced drop, and the type of opsin expressed, to assess their relative contributions to the post-illumination surge (Table 1). We found no correlation between the amplitude of the illumination-induced drop in [K^+^]_o_, and the rebound surge in [K^+^]_o_ for either experimental group individually (Table 1; Figure 3), whereas across the entire data set, virtually all the variance was explained by the type of opsin, and almost nothing by the size of the preceding, illumination-induced drop in [K^+^]_o_. This is consistent with our previous data showing that a large element of the post-illumination surge in halorhodopsin slices was sensitive to blockade of KCC2, indicating that the surge is boosted significantly by being coupled also to Cl^-^ extrusion in the halorhodopsin experiments.

**Table 1.** Linear regression model analyses of Archaerhodopsin and Halorhodopsin [K^+^]_extra_. Linear regression model analyses of the data presented in Figure 3, examining different models of the post-illumination rise in [K^+^]_extra_ predicted by the expression of either Halorhodopsin or ArchT, and the degree of [K^+^]_extra_ drop during the period of illumination. The 3^rd^ analysis (“[K^+^] drop (all)”) is of the dataset with ArchT and Halorhodopsin brain slices pooled together without distinction. This distinction is made in the 4^th^ analysis ((“[K^+^] drop, ArchT/Halo grouped”), whereas the 5^th^ analysis does not include information about the [K^+^] drop during illumination. Note that the adjusted R^2^ actually improves without the [K^+^] drop information, indicating that the post-illumination [K^+^] rise is explained almost solely by the difference between the opsins.

|  | R<sup>2</sup> | Adjusted R<sup>2</sup> | p value |
| --- | --- | --- | --- |
| [K<sup>+</sup>] rise vs [K<sup>+</sup>] drop (Halo) | 0.006 | -0.076 | 0.79 |
| [K<sup>+</sup>] drop (ArchT) | 0.099 | -0.001 | 0.35 |
| [K<sup>+</sup>] drop (all) | 0.554 | 0.535 | < 10<sup>-4</sup> |
| [K<sup>+</sup>] drop, ArchT/Halo grouped | 0.724 | 0.699 | < 10<sup>-6</sup> |
| ArchT/Halo grouped | 0.721 | 0.709 | < 10<sup>-7</sup> |

**Figure 3.**
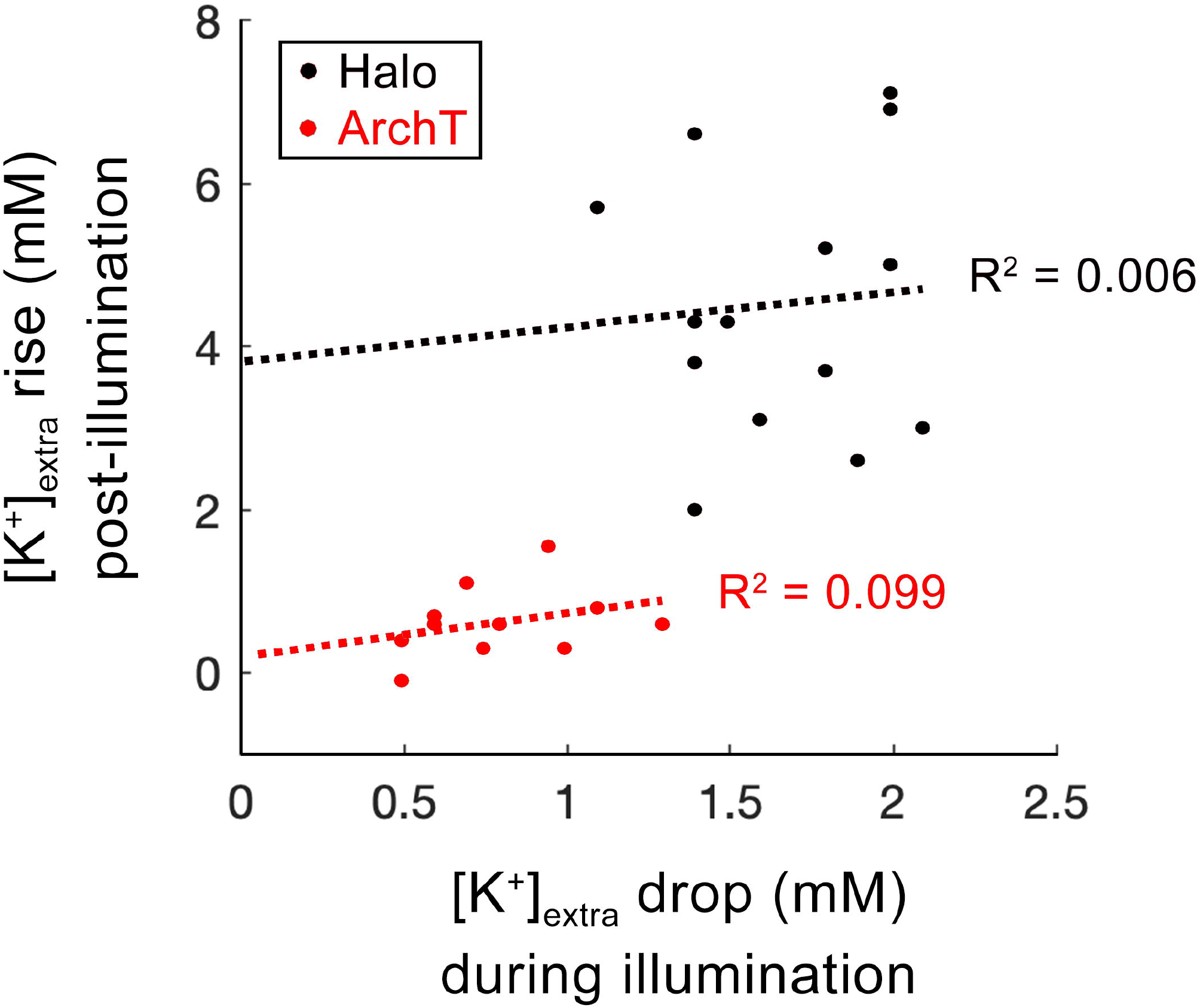
The post-illumination [K^+^]o rise does not reflect the drop in [K^+^]o during illumination. A plot of the [K^+^]_o_ rise, in excess of baseline (3.5mM), following the end of illumination versus the drop in [K^+^]o below baseline that occurs during illumination for 25 different brain slices, 14 expressing Halorhodopsin (red) and 11 expressing Archaerhodopsin-T (black). Halorhodopsin activation, using the expression systems available to us, produced a larger K^+^ dip during illumination (range 1.2-2.2mM below baseline) than did activation of ArchT (0.5-1.3mM below baseline). The dotted lines represent the correlation for the two experimental groups, extended to the ordinate axis. The difference in the intercepts represents the effect of chloride loading, secondary to halorhodopsin activation, producing an excess return of K^+^ to the extracellular space. Table 1 shows the R^2^ values for different linear regression models, which show that virtually all the experimental variation is explained in terms of the two experimental groups, and almost none by the intra-illumination [K^+^]_o_ drop.

A final point of difference is that in none of the archaerhodopsin experiments did we observe cortical spreading depolarization events; in contrast, these were triggered quite readily in halorhodopsin-expressing tissue (not illustrated, but examples are found in (Parrish et al., 2023)).

## Discussion

Archaerhodopsin and halorhodopsin have both been used to inhibit neuronal activity, but do so by driving ionic movement in opposite directions: halorhodopsin by driving Cl^-^ inwards, and archaerhodopsin by driving H^+^ out. We show here that this is further reflected by differences in the pattern of secondary effects, consistent with our interpretation of these effects reported in a previous study of halorhodopsin. The two optogenetic pumps share the effect of hyperpolarizing neurons, and consistent with this, both show a secondary drop in [K^+^]_o_, caused by K^+^ moving into the stimulated cells through leak channels driven by the altered electro-chemical balance for K^+^ during the period of opsin activity. The rebound, post-illumination surge in [K^+^]_o_, however, appeared much smaller for archaerhodopsin, even allowing for the fact that the expression levels of the two opsins were not exactly equivalent (halorhodopsin expression was achieved by cross-breeding Emx1 and floxed halorhodopsin mouse lines; archaerhodopsin expression was obtained by injecting Emx1 mice with a viral vector carrying the floxed archaerhodopsin gene) and so we were unable to achieve the same degree of K^+^ loading during the periods of illumination. Across the entire dataset, pooled from both halorhodopsin- and archaerhodopins-expressing brain slices, virtually all the variance in the peak amplitude of the rebound [K^+^]_o_ was explained by the choice of opsin, and very little by the within-illumination drop.

The much larger rebound surge in [K^+^]_o_ with halorhodopsin is, in fact, in line with what can be predicted from the different actions of the two opsin pumps. Halorhodopsin loads the cell with chloride, whereas archaerhodopsin consumes protons from the abundant intracellular buffer species. Even if halorhodopsin and archaerhodopsin current amplitudes and durations were identical, only halorhodopsin loads the cell with parallel inward fluxes of K^+^ and Cl^-^, leading to outward transport of KCl by KCC2, which is an electroneutral transporter capable of restoring a normal intraneuronal Cl^-^ level rapidly after a Cl^-^ load. This pattern of ionic redistribution is consistent with previous observations that the rebound [K^+^]_o_ surge, following chloride-loading arising either naturally, following intense GABAergic activation (Viitanen et al., 2010), or artificially from halorhodopsin activation (Parrish et al., 2023), is reduced by blockers of KCC2.

These collected observations, including the much smaller surge following archaerhodopsin activation, further reinforces the view that [Cl^-^]_i_ and [K^+^]_o_ are tightly coupled, as seen with the large rise in [K^+^]_o_ that follows intense interneuronal activation or direct GABA application (Chang et al., 2018; Viitanen et al., 2010). These observations warrant a reconsideration of the nature of raised [K^+^]_o_ during epileptic seizures, which may comfortably exceed 10mM at its peak (Hablitz and Heinemann, 1987; Librizzi et al., 2017; Raimondo et al., 2015; Somjen, 2004). This has often been attributed solely to K^+^ movement out of cells during action potentials, as described originally by Hodgkin and Huxley (Hodgkin and Huxley, 1952). Recordings of the ictal initiation period typically also show a significant rise in [K^+^]_o_ ahead of the time when pyramidal cells are recruited, as is readily apparent in published recordings (Chizhov et al., 2022; Librizzi et al., 2017). Notably, this early peak coincides with a period of very intense interneuronal activity (Parrish et al., 2019; Trevelyan et al., 2006). A major contribution to this early rise, therefore, is likely to be the extrusion of K^+^ coupled to Cl^-^-clearance, following a period of significant chloride-loading due to intense GABAergic bombardment. Recognising these two different causes of K^+^ extrusion – (1) the Hodgkin-Huxley model of action potential firing and (2) coupled to the clearance of Cl^-^ is important to our understanding of ictal propagation and recruitment to seizures (Graham et al., 2022; Trevelyan et al., 2022).

The most surprising secondary effect triggered by prolonged halorhodopsin activation was the occurrence of CSDs, at times, arising from a state where the majority of neurons were hyperpolarized to different degrees, and [K^+^]_o_ was below baseline levels (Parrish et al., 2023). We did not observe such events in our archaerhodopsin preparations. Two possible explanations suggest themselves for this difference, both consistent with our data: first that the lack of CSDs in our archaerhodopsin experiments reflects the smaller shifts in K^+^ (Figure 3), or second that it reflects differences in the net ion flux during illumination. Halorhodopsin activation results in both inward Cl^-^ and K^+^ movement, generating a large net influx of osmotically active particles. In contrast, archaerhodopsin drives an outward proton movement, and the K^+^ redistribution is much smaller. While this negative finding (no CSDs in our archaerhodopsin experiments) represents a limited test of the hypothesized differences between halorhodopsin and archaerhodopsin, the result is consistent with what else is known about their different mechanisms of action.

## Acknowledgments

The work was supported by grants from BBSRC (BB/P019854/1) and MRC (MR/R005427/1). We would like to thank the support staff in the animal facilities at Newcastle University.

